# High glucose-induced ubiquitylation of G6PD leads to the injury of podocyte

**DOI:** 10.1101/350694

**Authors:** Meng Wang, Ji Hu, Linling Yan, Yeping Yang, Min He, Shizhe Guo, Meng Wu, Qin Li, Wei Gong, Yang Yang, Diane E. Handy, Bin Lu, Chuanming Hao, Qinghua Wang, Yiming Li, Ronggui Hu, Robert C. Stanton, Zhaoyun Zhang

## Abstract

Oxidative stress contributes substantially to podocyte injury in diabetic kidney disease. The mechanism of hyperglycemia-induced oxidative stress in podocytes is not fully understood. Glucose-6-phosphate dehydrogenase is critical in maintaining NADPH, an important cofactor for antioxidant system. Here, we hypothesized that high glucose induces ubiquitylation and degradation of G6PD, which injures podocytes by reactive oxygen species (ROS) accumulation. We found that both G6PD protein expression and G6PD activity was decreased in kidneys of both diabetic patients and diabetic rodents. Overexpressing G6PD reversed redox imbalance and podocyte apoptosis induced by high glucose and palmitate. Inhibition of G6PD induced podocyte apoptosis. In G6PD deficient mice, podocyte apoptosis was also largely increased. High glucose had no effect on G6PD mRNA level but it caused decreased G6PD protein expression, which was mediated by the ubiquitin proteasome pathway. Furthermore, von Hippel−Lindau (VHL), an E3 ubiquitin ligase subunit, directly bound to G6PD and degraded G6PD through ubiquitylating G6PD on lysine residues 366/403. Our data suggest that high glucose induces ubiquitylation of G6PD by VHL, which leads to ROS accumulation and podocyte injury.

## Introduction

Diabetic kidney disease (DKD) is the major cause of end-stage renal disease. Chronic hyperglycemia leads to the injury and dysfunction of podocytes, which plays an important role in the development and progression of DKD[1–3]. Due to the key role of podocytes in maintaining the glomerular filtration, recent studies have been focused on protecting against podocyte injury and loss in various glomerular diseases, including DKD. Currently, there are several agents under clinical investigation for the treatment of DKD targeting podocyte [4]. However, the underlying mechanisms as well as the pathogenesis of podocyte injury and loss remain largely unknown[5–8].

Recent studies have suggested that increased oxidative stress plays an important role in the podocyte injury and loss of DKD[9]. The accumulation of reactive oxidative species (ROS) is caused by the imbalance between processes that produce ROS and processes that reduce ROS. The antioxidant system is consisted of catalase, superoxide dismutase, glutathione system and thioredoxin system[10, 11]. For both glutathione and thioredoxin systems, nicotinamide adenine dinucleotide phosphate (NADPH) is the required cofactor for the conversion of oxidized glutathione and thioredoxin to the reduced forms, which scavenge ROS. In addition, there is an allosteric binding site for NADPH in catalase. The binding of NADPH maintains catalase in its most active tetrameric conformation and protects it against the toxicity of hydrogen peroxide[12]. Thus, NADPH is a critical component for the antioxidant system.

Our and others’ previous studies suggest that the pentose phosphate pathway (PPP) is the principal pathway for producing NADPH, in which glucose 6-phosphate dehydrogenase (G6PD) is the rate-limiting enzyme[13–16]. We and others have shown that high glucose decreases G6PD activity in endothelial cells, pancreatic β cells, kidney tissue, liver tissue, and pancreas tissue, which leads to insufficient NADPH supply and thus the accumulation of ROS[16–21]. We also found that G6PD deficient mice had increased renal oxidative stress and elevated urinary albumin, which suggested that G6PD deficiency alone was sufficient to injure the glomerular filtration barrier[22]. However, the mechanism of high glucose-mediated decrease in G6PD activity is unknown. To address this important question, we investigated the regulatory mechanism(s) that affected G6PD in podocytes under hyperglycemia. Our findings suggest that hyperglycemia-induced ubiquitylation of G6PD is a major contributor to the injury and loss of podocytes, which might be a drug target for DKD treatment.

## Results

### G6PD protein expression and activity was decreased in diabetic kidney

In order to examine the expression of G6PD protein in diabetes mellitus, first we checked the renal cortex from non-diabetic subjects (n=3) and diabetic patients (n=3) by immunohistochemical (IHC) staining. Compared to non-diabetic subjects, the renal G6PD protein expression was decreased in diabetic patients (Figure 1A). Next, we explored G6PD level in different diabetic rodents including STZ-induced diabetic rats, STZ-induced diabetic mice and Akita mice (a model of type 1 diabetes). Compared to non-diabetic (NDM) controls, blood glucose was significantly increased in diabetic rodents (Figure EV1A, EV1B and EV1C). Meanwhile, G6PD protein expression in renal cortex was largely decreased in diabetic models (Figure 1B, 1C and 1D). Further, G6PD activity was examined. As shown in Figure 1E, diabetic mice have lower G6PD activity in the renal cortex compared to NDM controls.

**Figure 1.**
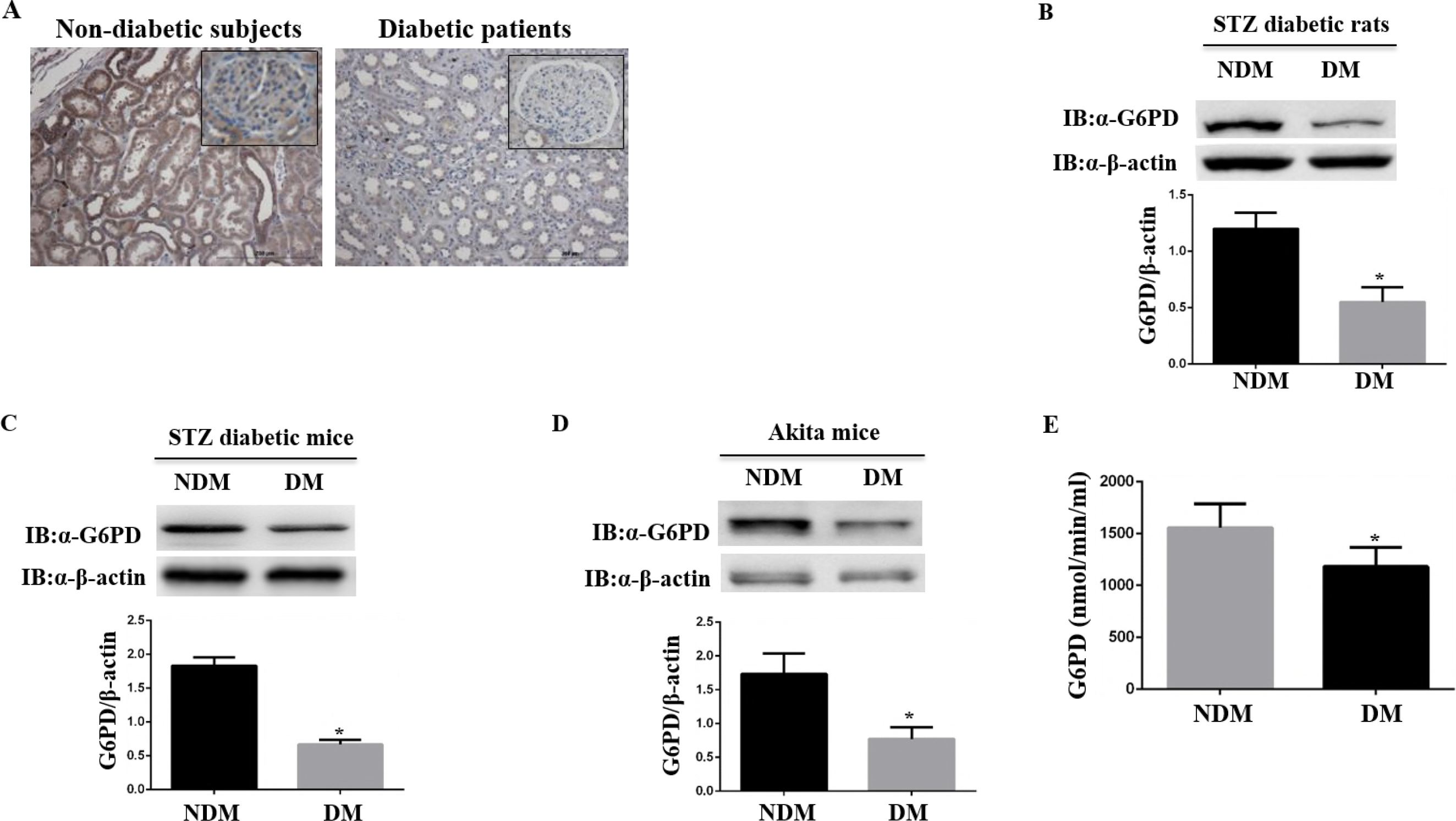
G6PD protein expression and activity were decreased in diabetic kidney. A G6PD protein expression was significantly decreased in diabetic kidney. The renal cortex from non-diabetic subjects (n=3) and diabetic patients (n=3) were examined by IHC staining for G6PD. Shown are average values with standard deviation (s.d.). ^****^ denotes *P* < 0.0001 for DM versus NDM. B, C, D G6PD protein level was decreased in different diabetic rodents, including STZ diabetic rats (B), STZ diabetic mice (C) and Akita mice (D). The renal cortex from non-diabetic (NDM) controls and diabetic rodents (DM) were collected and Western blot was performed to examine the expression of G6PD protein. Shown are average values with standard deviation (s.d.). n=5 mice for each group. ^*^ denotes *P* < 0.05 for DM versus NDM. E G6PD activity was decreased in the diabetic kidney. The renal cortex from non-diabetic (NDM) controls and STZ-induced diabetic mice (DM) were collected and G6PD activity was determined. Shown are average values with standard deviation (s.d.). n=5 mice for each group. ^*^ denotes *P* < 0.05 for DM versus NDM.

### G6PD expression was decreased in podocytes of diabetic kidney

Podocytes line the outer aspect of the glomerular basement membrane (GBM) and are highly differentiated. Podocyte injury and loss indicates it’s associated with the initiation and development of DKD. Whether low expression of G6PD is associated with podocyte injury and loss in DKD has not been illuminated. Wilms’ tumor protein-1 (WT-1) is a specific marker for podocytes. To examine the podocyte number, immunofluorescence against WT-1 was performed. As shown in Figure 2A, compared to the NDM, the podocyte number, reflected by WT-1 positive staining, in renal cortex of DM was fewer. To examine the G6PD protein expression in podocytes in vivo, immunofluorescence co-staining of WT-1 and G6PD was performed in the kidney. Compared to NDM, G6PD protein expression in podocytes of diabetic mice was decreased (Figure 2B).

**Figure 2.**
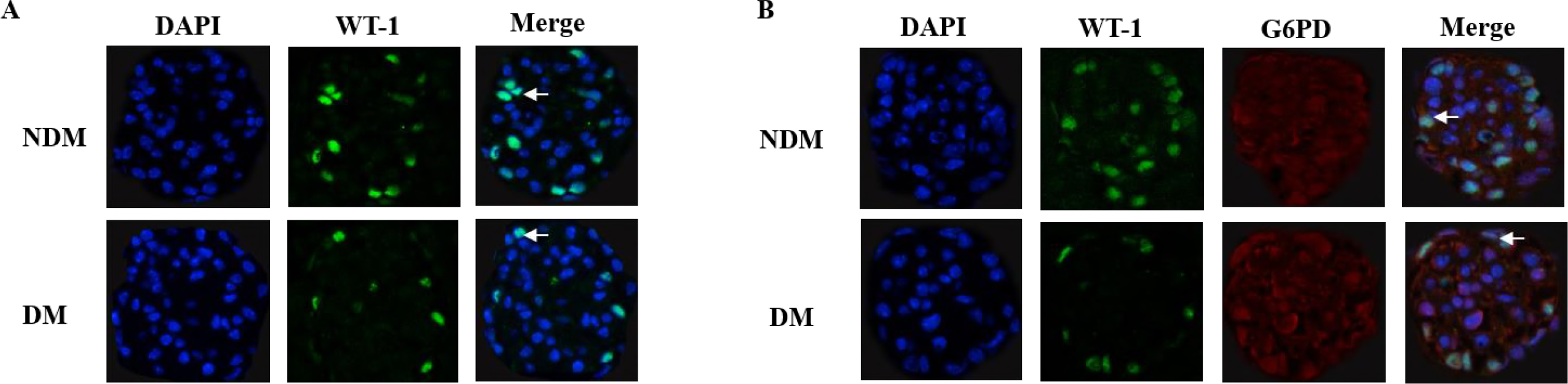
G6PD expression was decreased in podocytes of diabetic kidney. A Podocyte number was decreased in diabetic kidney. The renal cortex from non-diabetic (NDM) controls and diabetic rodents (DM) were examined with immunofluorescence using anti-WT-1 antibody to label podocyte cells (green staining). The nuclei were counterstained with 4′,6-diamidino-2-phenylindole (DAPI, blue staining). n=4 mice for each group. Magnification 40×. B G6PD protein level was decreased in podocytes of diabetic kidney. The renal cortex from non-diabetic (NDM) controls and diabetic rodents (DM) were examined with co-immunofluorescence using anti-WT-1 antibody (green staining) and anti-G6PD antibody (red staining). DAPI was used to label the nuclei (blue staining). Colocalization of the fluorochromes yielded a yellow color (see arrows). n=4 mice for each group. Magnification 40×.

### High glucose and palmitate decreased G6PD protein expression and increased apoptosis of podocyte

Previous studies have shown that both high glucose and lipids such as palmitate could induce podocyte injury[23–25]. As shown in Figure 2, renal G6PD protein expression declined in podocytes, while the causal relationship with metabolic disturbances was unknown. We conducted in vitro studies in cultured podocytes using either high glucose or palmitate. Mannitol (M) was added as an osmotic control. Compared to cells cultured in normal glucose (5.6mM glucose), G6PD protein expression of podocytes in high glucose (25mM glucose) was significantly decreased (Figure 3A). Meanwhile, increased apoptosis of podocytes was observed as reflected by increased expression of cleaved caspase-3 (Figure 3A). Similarly, palmitate also reduced the protein expression of G6PD and induced apoptosis of podocytes in a dose-dependent manner (Figure 3B).

**Figure 3.**
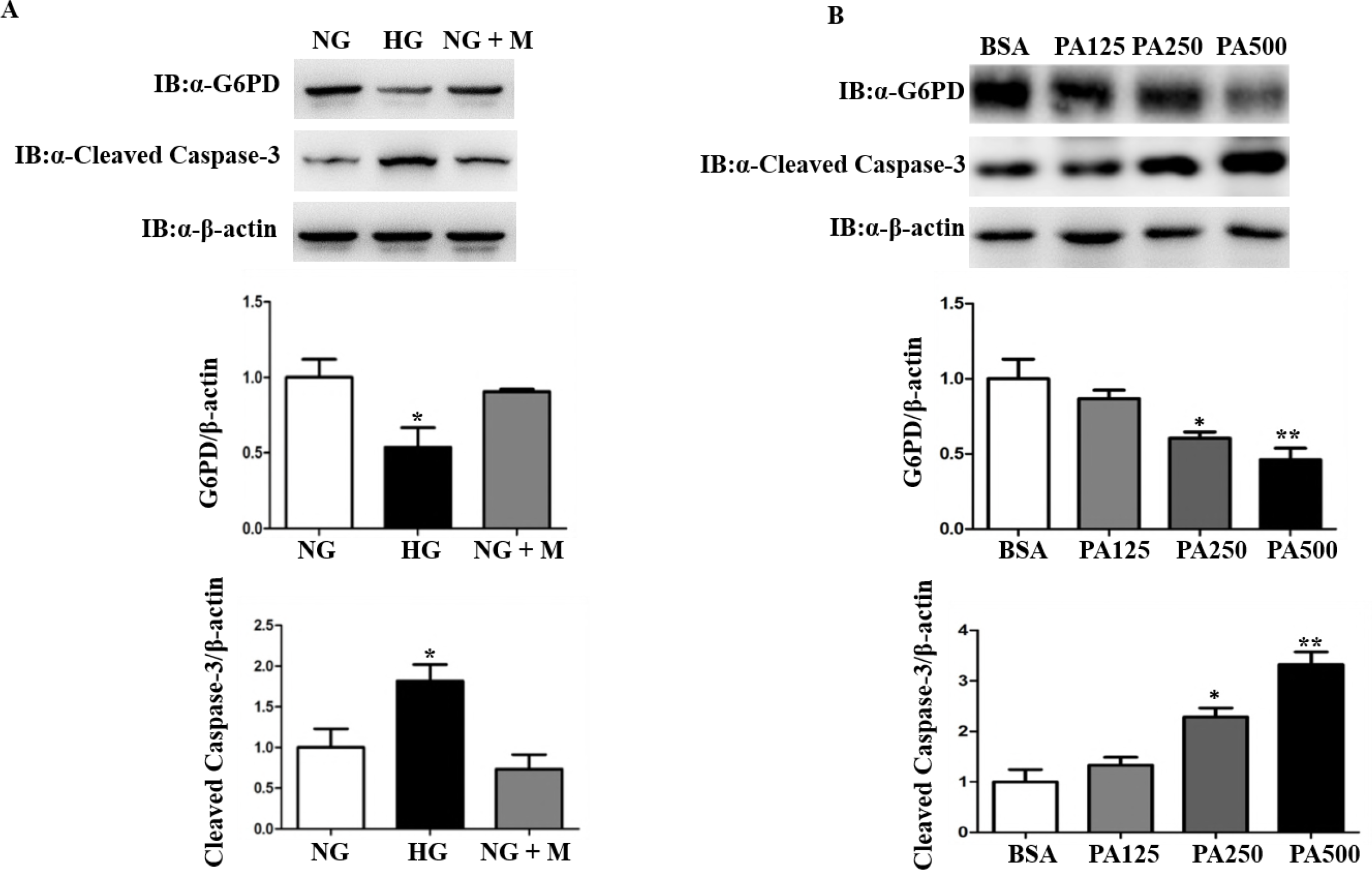
High glucose and palmitate decreased G6PD protein expression and increased apoptosis of podocyte. A High glucose decreased G6PD protein level and increased podocytes apoptosis. Podocytes were treated with normal glucose (NG, 5.6mM glucose), high glucose (HG, 25mM glucose), or normal glucose supplemented with 19.4mM mannitol (NG+M) for 72 hours. Mannitol (M) was added as an osmotic control. The protein levels of G6PD and cleaved caspase-3 were determined by Western blot. Shown are average values with standard deviation (s.d.) of triplicated experiments. ^*^ denotes *P* < 0.05 for cells treated with HG versus cells with NG or NG + M. B Palmitate decreased the expression of G6PD protein and increased the apoptosis of podocyte. Podocytes were treated with bovine serum albumin (BSA), 125μM palmitate (PA125), 250μM palmitate (PA250) or 500μM palmitate (PA500) for 24 hours. Protein levels of G6PD and cleaved caspase-3 were determined by Western blot. Shown are average values with standard deviation (s.d.) of triplicated experiments. ^*^ denotes *P* < 0.05 for cells treated with PA250 versus cells with BSA and ^**^ denotes *P* < 0.01 for cells treated with PA500 versus cells with BSA.

### Inhibition of G6PD increased apoptosis and loss of podocytes both in vitro and in vivo

To elucidate whether G6PD deficiency per se affected podocytes survival, siRNA targeting to G6PD (siG6PD) was constructed to inhibit the expression of G6PD. Mouse podocytes were transfected with either siG6PD or scrambled siRNA (scramble) as a control (Figure 4A). Inhibition of G6PD increased apoptosis of podocytes (Figure 4B). To further determine if G6PD deficiency per se would affect the survival of podocytes in vivo, hemizygous (Hemi) G6PD deficient mice were employed in this study. Previous studies showed that G6PD activity in Hemi G6PD deficient mice dramatically decreased (by 85%), as compared to the control mice[22, 26]. In order to clarify the indispensable role of G6PD in podocytes survival, renal cortex from Hemi G6PD deficient mice at different ages and age-matched littermate wild type mice were examined with IHC for WT-1. Compared to age-matched wild type mice, G6PD deficient mice at 9-week-old (9wk), 23-week-old (23wk) and 39-week-old (39wk) all showed decreased podocyte number in kidney (Figure 4C). Interestingly, as compared to 9wk G6PD deficient mice, 39wk G6PD deficient mice showed even fewer podocytes (Figure 4C), suggesting that longer duration of G6PD deficiency caused more podocytes loss. Further, immunofluorescence co-staining for both WT-1 and TUNEL assay was performed to evaluate the apoptosis of podocytes. Compared to age-matched wild type mice, TUNEL positive cells were increased in the Hemi G6PD deficient mice (Figure 4D).

**Figure 4.**
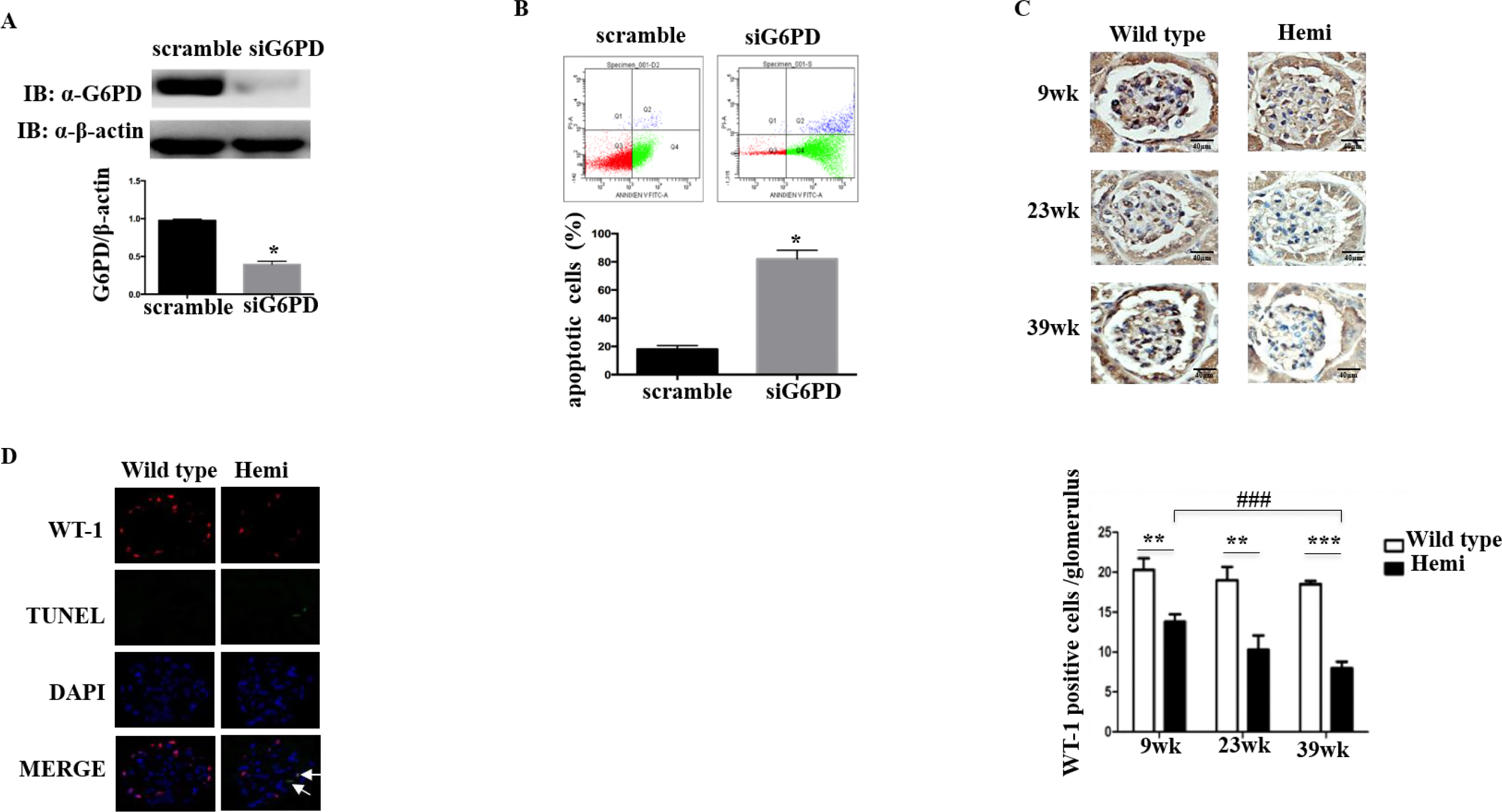
Inhibition of G6PD increased apoptosis and loss of podocytes both in vitro and in vivo. A siRNA targeting to G6PD (siG6PD) significantly inhibited G6PD protein level. Podocytes were transfected with either siG6PD or scrambled siRNA for 48 hours and then G6PD protein expression was determined by Western blot. Shown are average values with standard deviation (s.d.) of triplicated experiments. ^*^ denotes *P* < 0.0′5 for cells transfected with siG6PD versus cells with scramble. B Inhibition of G6PD led to the increased apoptosis of podocyte. Podocytes were transfected with either scrambled siRNA (scramble) or siG6PD and flow cytometry was used to detect podocytes apoptosis. Shown are average values with standard deviation (s.d.) of triplicated experiments. ^*^ denotes *P* < 0.05 for cells transfected with siG6PD versus cells with scramble. C Deficiency of G6PD caused podocytes loss. The renal cortex from Hemi G6PD deficient mice aged 9wk, 23wk and 39wk and age-matched littermate wild type mice were examined with IHC staining for WT-1. n=6 mice for each group. Magnification 40×. ** denotes *P* < 0.01 and ^***^ denotes *P* < 0.001 for Hemi G6PD deficient mice versus wild type mice. ### denotes *P* < 0.001 for 9wk Hemi G6PD deficient mice versus 39wk Hemi G6PD deficient mice. D Deficiency of G6PD caused increased podocyte apoptosis in vivo. The renal cortex from Hemizygous (Hemi) G6PD deficient mice (right panel) and age-matched littermate wild type mice (left panel) were examined with co-immunofluorescence using anti-WT-1 antibody to label podocyte cells (red staining) and TUNEL assay to label apoptotic cells (green staining). The nuclei were counterstained with DAPI (blue staining). Colocalization of the fluorochromes results in a yellow color (see arrows). n=6 mice for each group. Magnification 40×.

### Elevation of G6PD expression decreased the apoptosis of podocyte

If high glucose or palmitate-induced decrease of G6PD protein plays a role in the survival of podocyte, overexpression of G6PD should reverse the apoptosis. To test this, podocytes were transfected with adenoviral G6PD vector (Ad-G6PD) to overexpress G6PD and apoptosis was examined. Empty vector adenoviruses (Ad-null) were used as a control. As shown in Figure 5, elevation of G6PD protein significantly decreased podocyte apoptosis induced by either high-glucose (Figure 5A) or palmitate (Figure 5B).

**Figure 5.**
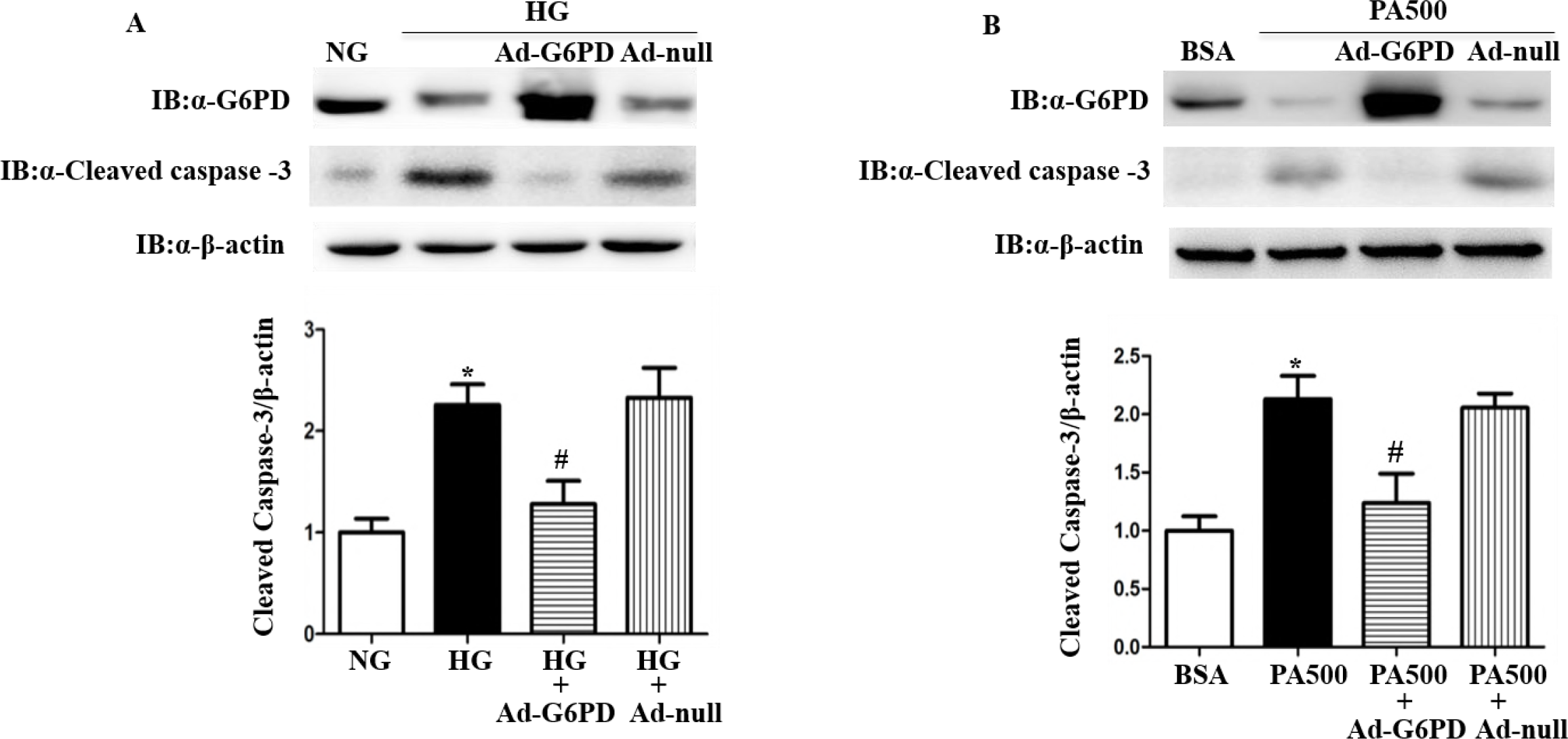
Elevation of G6PD expression decreased the apoptosis of podocyte. A Overexpressing G6PD ameliorated podocytes apoptosis caused by high glucose. Podocytes were treated with normal glucose (NG, 5.6mM glucose), high glucose (HG, 25mM glucose), high glucose with infection of adenoviruses G6PD (HG+Ad-G6PD) and high glucose with infection of empty vector adenoviruses (HG+Ad-null). The levels of protein were determined by Western blot. Shown are average values with standard deviation (s.d.) of triplicated experiments. ^*^ denotes *P* < 0.05 for cells treated with HG versus cells with NG. # denotes *P* < 0.05 for cells treated with HG+Ad-G6PD versus cells with HG+Ad-null. B Elevation of G6PD rescued the apoptosis of podocyte induced by palmitate. Podocytes were treated with bovine serum albumin (BSA), 500μM palmitate (PA500), 500μM palmitate with infection of adenoviruses G6PD (PA500+Ad-G6PD) and 500μM palmitate with infection of empty vector adenoviruses (PA500+Ad-null). Protein expression were determined by Western blot. Shown are average values with standard deviation (s.d.) of triplicated experiments. ^*^ denotes *P* < 0.05 for cells treated with PA500 versus cells with BSA. # denotes *P* < 0.05 for cells treated with PA500+Ad-G6PD versus cells with PA500+Ad-null.

### Elevation of G6PD protein expression reversed the redox imbalance caused by high glucose

High glucose-induced increase of ROS in podocytes has been validated in previous studies[27, 28]. In order to elucidate the role of G6PD in high glucose-induced oxidative stress, we overexpressed G6PD in podocytes exposed to high glucose and examined parameters reflecting oxidative stress. Empty vector adenoviruses (Ad-null) were used as control. Compared to cells cultured in normal glucose, NADPH level in podocytes exposed to high glucose was significantly decreased, which was reversed by overexpressing G6PD (Figure 6A). NADPH is required for oxidized glutathione (GSSG) to be converted to reduced glutathione (GSH), a major component in antioxidant system. Thus, we checked the level of GSH/GSSG of podocytes and found that high glucose decreased GSH/GSSG, which was increased by G6PD overexpression (Figure 6B). As a consequence, high glucose-induced ROS accumulation in podocytes was found to be decreased by overexpressing G6PD (Figure 6C).

**Figure 6.**
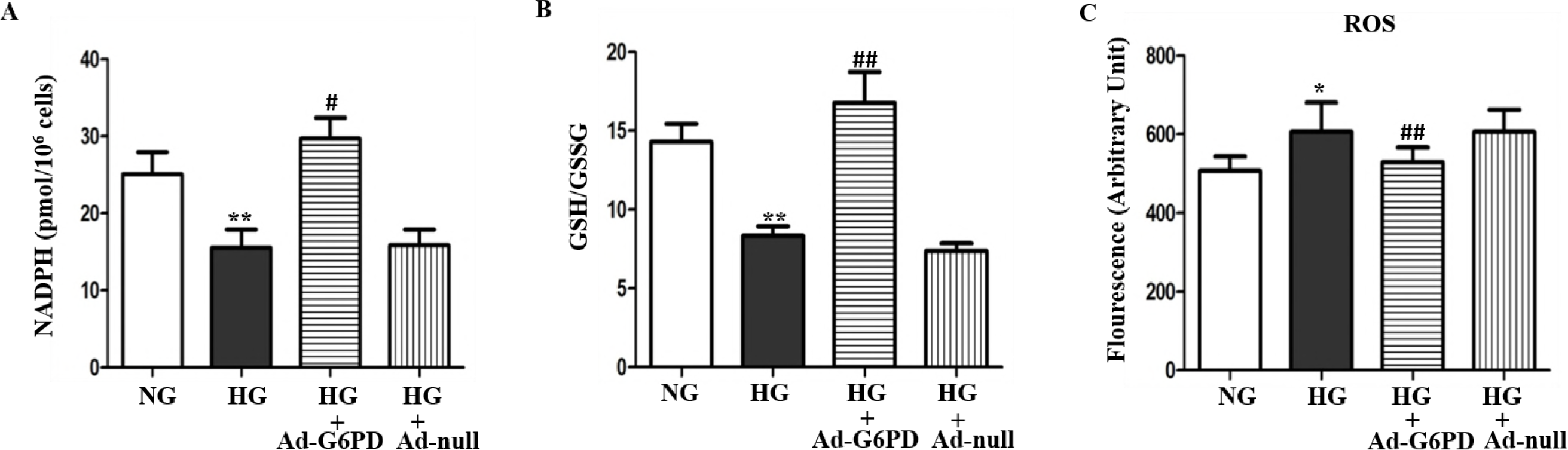
Elevation of G6PD protein expression reversed the redox imbalance caused by high glucose. A Overexpressing G6PD increased NADPH level in podocytes exposed to high glucose. Podocytes were treated with normal glucose (NG, 5.6mM glucose), high glucose (HG, 25mM glucose), high glucose with infection of adenoviruses G6PD (HG+Ad-G6PD) and high glucose with infection of empty vector adenoviruses (HG+Ad-null). NADPH was measured by a colorimetric method according to the manufacturer’s instructions. Shown are average values with standard deviation (s.d.) of triplicated experiments. ^**^ denotes *P* < 0.01 for cells treated with HG versus cells with NG. # denotes *P* < 0.05 for cells treated with HG+Ad-G6PD versus cells with HG+Ad-null. B the decreased GSH/GSSG in podocytes exposed to high glucose was ameliorated by overexpressing G6PD. Podocytes were treated as described in (A) and GSH/GSSG was measured by a spectrophotometric method following the manufacturer’s instructions. Shown are average values with standard deviation (s.d.) of triplicated experiments. ^**^ denotes *P* < 0.01 for cells treated with HG versus cells with NG. ## denotes *P* < 0.01 for cells treated with HG+Ad-G6PD versus cells with HG+Ad-null. C Elevation of G6PD protein expression reduced the accumulation of ROS in podocytes exposed to high glucose. Podocytes were treated as described in Figure 6A. ROS accumulation was measured with the cell-permeable, oxidation-sensitive dye CM-H_2_DCFDA. Shown are average values with standard deviation (s.d.) of triplicated experiments. ^*^ denotes *P* < 0.05 for cells treated with HG versus cells with NG. ## denotes *P* < 0.01 for cells treated with HG+Ad-G6PD versus cells with HG+Ad-null.

### High glucose promoted G6PD protein degradation through the ubiquitin proteasome pathway

To investigate the mechanism underlying high glucose or palmitate-induced decrease in G6PD protein level, we examined the G6PD mRNA level in podocytes under either high glucose or palmitate incubation condition. Notably, palmitate significantly decreased G6PD mRNA level (Figure 7A), while high glucose had no effect on it (Figure 7B). Furthermore, G6PD protein level was found to be decreased in podocytes exposed to high glucose for 72 hours (Figure 7C). Thus, the reduction of G6PD protein by high glucose was unlikely to be due to transcriptional regulation and might be associated with the protein degradation. As the ubiquitin (Ub) proteasome pathway (UPP) degraded the majority of intracellular proteins, we questioned whether this pathway was involved in decreasing G6PD protein induced by high glucose. To this end, MG132, a proteasome inhibitor, was added, which significantly rescued G6PD protein level in podocytes exposed to high glucose (Figure 7D). To confirm the ubiquitylation of G6PD, Flag-tagged G6PD (Flag-G6PD) and His-tagged ubiquitin (His-Ub) plasmids were constructed and validated in HEK293T cells (Figure EV2A and EV2B). Subsequently, Flag-G6PD and His-Ub constructs were co-transfected into HEK293T cells and anti-Flag M2 beads were used for immunoprecipitation (IP). Western blot analysis using anti-His antibody showed that G6PD was indeed ubiquitylated and the ubiquitylation was largely enhanced in the presence of MG132 (Figure 7E). These results indicated that high glucose-induced G6PD protein decrease depended on the ubiquitin proteasome pathway.

**Figure 7.**
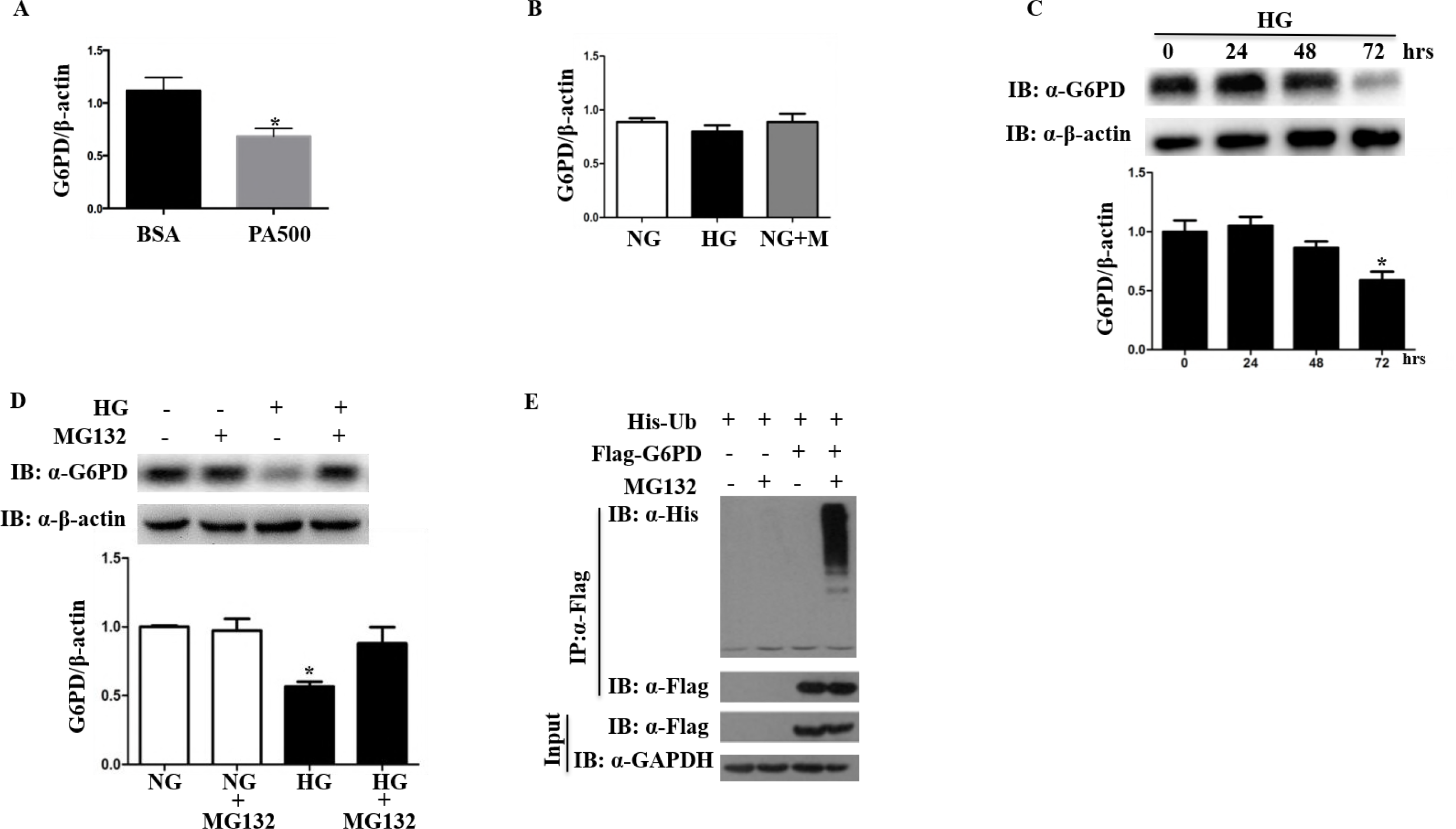
High glucose promoted G6PD protein degradation through the ubiquitin proteasome pathway. A Palmitate decreased the level of G6PD mRNA. Podocytes were treated with BSA, 500μM palmitate (PA500) for 24 hours. G6PD mRNA level was measured by real-time PCR. Shown are average values with standard deviation (s.d.) of triplicated experiments. ^*^ denotes *P* < 0.05 for cells treated with PA500 versus cells with BSA. B High glucose had no effect on G6PD mRNA level. Podocytes were treated with normal glucose (NG, 5.6mM glucose), high glucose (HG, 25mM glucose), or normal glucose supplemented with mannitol (NG+M) for 72 hours. Real-time PCR was used to analyze G6PD mRNA abundance. Shown are average values with standard deviation (s.d.) of triplicated experiments. C The degradation of G6PD induced by high glucose was in a time-dependent manner. Podocytes were incubated with high glucose (HG, 25mM glucose) for the indicated time, and cell lysates were subjected to G6PD and β-actin immunoblotting. Shown are average values with standard deviation (s.d.) of triplicated experiments. ^*^ denotes *P* < 0.05 for cells treated with HG for 72 hours versus cells incubated in normal glucose. D The decreased G6PD protein level in high glucose was largely rescued by MG132. Podocytes were cultured with normal glucose (NG, 5.6mM glucose), normal glucose supplemented with 0.5μM MG132 (NG+MG132), high glucose (HG, 25mM glucose) or high glucose supplemented with 0.5μM MG132 (HG+MG132). Shown are average values with standard deviation (s.d.) of triplicated experiments. ^*^ denotes *P* < 0.05 for cells treated with HG versus cells with HG+MG132. E G6PD was ubiquitylated. HEK293T cells were transfected with indicated plasmids with or without 20μM MG132 for 12 hours. Cell lysates were subjected to immunoprecipitation (IP) with anti-Flag M2 beads. The precipitates were probed using His and Flag antibodies. Input cell lysates were subjected to Flag and GAPDH immunoblotting.

### VHL ubiquitylated and degraded G6PD

The E3 ubiquitin ligases are the major enzymes responsible for recognizing and linking ubiquitin to the target proteins. To identify the potential E3 ligase that ubiquitylated G6PD, we conducted yeast two hybrid (Y2H) screening[29]. Specifically, human G6PD was employed as the bait to screen potential G6PD-interacting proteins from a Y2H prey library containing open reading frames from human cDNAs encoding over 400 putative ubiquitin ligases or their substrate binding subunits. We found that von Hippel-Lindau (VHL), a subunit of E3 ubiquitin ligase, interacted with G6PD in the Y2H system (Figure 8A). A VHL plasmid was constructed with an HA tag (HA-VHL) and was validated in HEK293T cells (Figure EV2C). The specific interaction between G6PD and VHL was verified with co-immunoprecipitation (CO-IP) assay. As shown in Figure 8B, Flag-G6PD and HA-VHL proteins formed a complex that survived the multistep procedures in CO-IP assay. Collectively, the specific interaction between VHL and G6PD was verified in the yeast two hybrid and CO-IP assay in cultured HEK293T cells, indicating that VHL was a G6PD-interacting protein.

**Figure 8.**
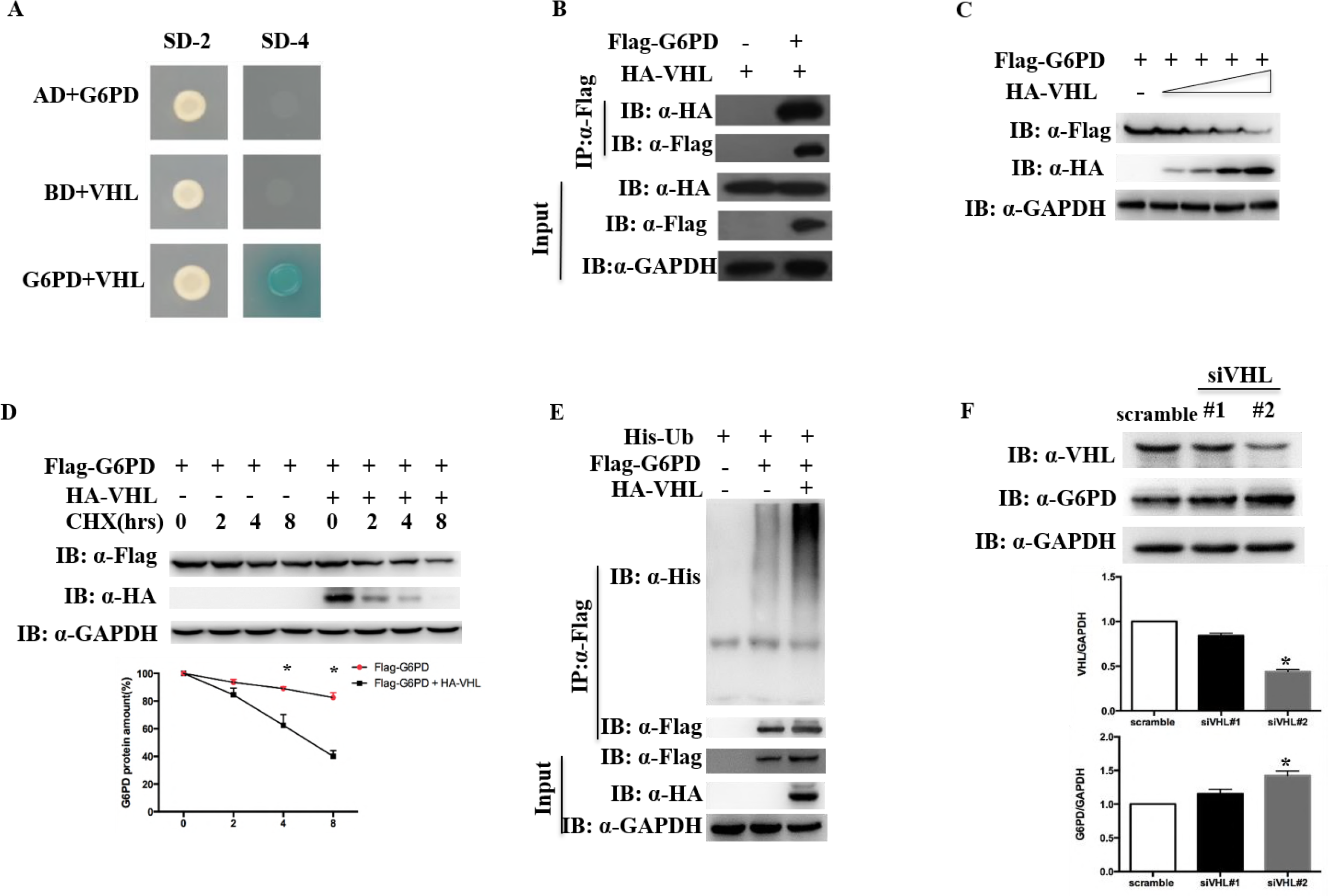
VHL ubiquitylated and degraded G6PD. A Human G6PD interacted with the E3 ubiquitin ligase VHL in Y2H system. Using human G6PD as bait, VHL was identified as an interacting protein with G6PD in yeast. G6PD and VHL were co-transformed into yeast strain Mav203-activated expression of β-galactosidase. AD, Activation Domain; BD, Binding Domain; SD-2, deficient in Leucine and Tryptophan; SD-4, deficient in Leucine, Tryptophan, Histidine, and Uracil. B G6PD interacted with VHL. Co-immunoprecipitation assay shown that tagged G6PD and VHL formed a complex in HEK293T cells. HEK293T cells were transfected with HA-VHL expression plasmid in combination with or without Flag-G6PD expression plasmid. Flag-G6PD was immunoprecipitated with anti-Flag M2 beads. The precipitates were probed using HA and Flag antibodies. C VHL negatively regulated the expression of G6PD protein. HEK293T cells were co-transfected with Flag-G6PD expression plasmid and increasing amounts of HA-VHL expression plasmid. The levels of Flag and HA were determined by Western blot with indicated antibodies. D VHL reduced G6PD protein stability. HEK293T cells were transfected with Flag-G6PD expression plasmid with or without HA-VHL expression plasmid. After 36 hours, cells were treated with CHX (100μg/ml) for the indicated time. Cell lysates were subjected to Flag, GAPDH and HA immunoblotting. Shown are average values with standard deviation (s.d.) of triplicated experiments. ^*^ denotes *P* < 0.05 for cells co-transfected with Flag-G6PD and HA-VHL (Flag-G6PD + HA-VHL) versus cells transfected with Flag-G6PD. E G6PD was efficiently ubiquitylated in the presence of VHL. HEK293T cells were transfected with different combinations of plasmids as indicated. G6PD was immunoprecipitated with anti-Flag M2 beads and immunoblotted with anti-His antibody to detect ubiquitylated G6PD. F Knockdown of endogenous VHL enhanced G6PD protein abundance. HEK293T cells were transfected with scrambled siRNA (scramble), siVHL#1 and siVHL#2. The levels of endogenous G6PD and VHL were analyzed by Western blot with indicated antibodies. Shown are average values with standard deviation (s.d.) of triplicated experiments. ^*^ denotes *P* < 0.05 for cells treated with siVHL#2 versus cells with scramble.

To further examine whether VHL functioned as an E3 ubiquitin ligase against G6PD, the influence of VHL on G6PD protein abundance was determined. As shown in Figure 8C, G6PD protein level declined with the increased VHL protein expression. To further confirm the post-translational regulation of VHL on G6PD, cycloheximide (CHX), an inhibitor of protein synthesis, was used. As shown in Figure 8D, with the inhibition of protein synthesis by CHX, VHL also reduced G6PD protein stability. To further ascertain the functional interaction of VHL with G6PD, ubiquitylation of G6PD by VHL was explored under in vivo conditions. As shown in Figure 8E, G6PD was conspicuously polyubiquitylated when HA-VHL was expressed along with His-Ub. Furthermore, we assessed the effect of VHL knockdown on the G6PD protein. First, two siRNAs targeting to VHL (siVHL#1 and siVHL#2) were constructed. HEK293T cells were transfected with scrambled siRNA (scramble), siVHL#1 or siVHL#2 and G6PD protein level was analyzed. As shown in Figure 8F, endogenous VHL was silenced efficiently by siVHL#2. The knockdown of VHL concomitantly resulted in the elevation of endogenous G6PD protein level (Figure 8F). These results suggested that VHL functioned as an E3 ubiquitin ligase against G6PD.

### K^366^ and K^403^ in G6PD were the major sites for VHL-mediated ubiquitylation

After an in vivo ubiquitylation reaction, ubiquitylated G6PD was enriched with anti-Flag M2 beads and subjected to tryptic digestion, followed by mass spectra (MS) analysis. The attachment of ubiquitin to the side chain of a lysine (Lys) residue renders it resistant to trypsin cleavage, and tryptic digestion of ubiquitin (chains) attached to the site leaves a-Gly-Gly-group, originating from the C terminus of ubiquitin, on the side chain of the modified Lys[30]. Based on these signature features, ubiquitylation sites in G6PD were identified from tandem mass spectrometry (MS/MS) spectra. Altogether, we found that a total of 5 Lys residues (K^97^, K^366^, K^403^, K^407^ and K^408^) were ubiquitylated by VHL (Figure 9A).

**Figure 9.**
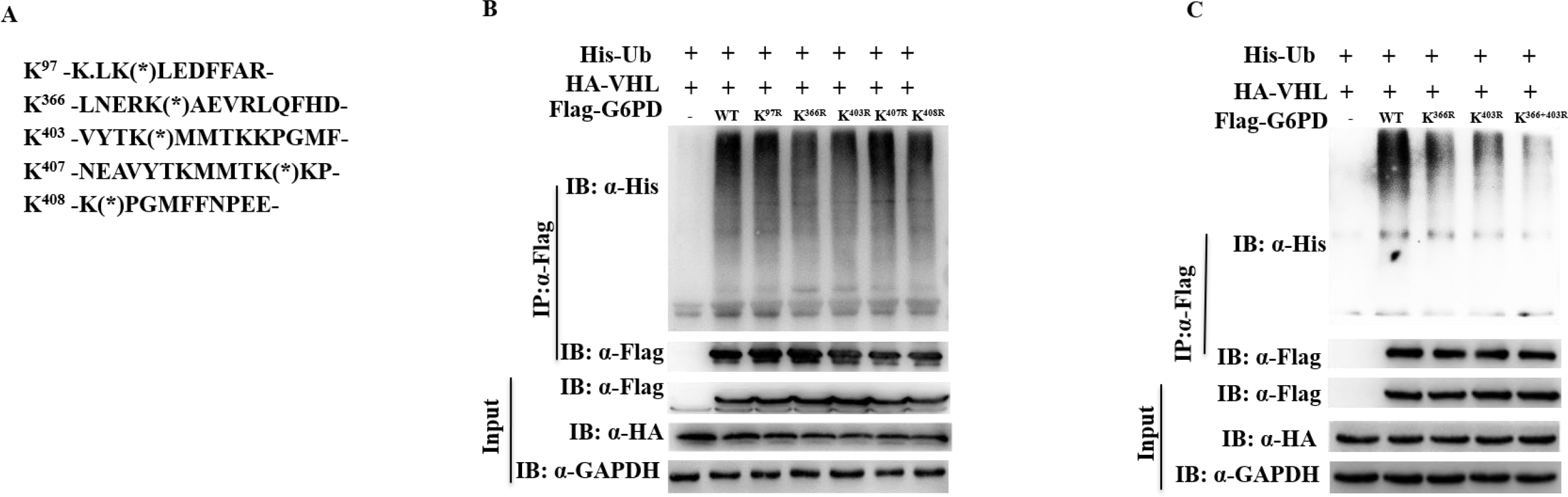
K^366^ and K^403^ in G6PD were the major sites for VHL-mediated ubiquitylation. A The map of the 5 lysine sites for VHL mediated ubiquitylation on G6PD. HEK293T cells were transfected with Flag-G6PD, His-Ub and HA-VHL plasmids, and then ubiquitylated G6PD were immunoprecipitated with anti-Flag M2 beads. MS spectra analysis identified 5 Lys residues (K^97^, K^366^, K^403^, K^407^and K^408^) for VHL-mediated ubiquitylation in G6PD. B K^366^ and K^403^ were the major sites for VHL-mediated ubiquitylation on G6PD. HEK293T cells were transfected with Flag-tagged wild-type G6PD (WT) or its mutants bearing single Lys-to-Arg substitutions at the above 5 potential ubiquitylation sites. Cell lysates were subjected to immunoprecipitation (IP) with anti-Flag M2 beads. The precipitates were probed using His and Flag antibodies. C Mutants with both K^366^ and K^403^ sites (K^366^+^403R^) largely abolished the ubiquitylation of G6PD. HEK293T cells were transfected with Flag-tagged wild-type G6PD (WT) or G6PD mutants bearing single or two Lys-to-Arg substitutions at K^366^ and K^403^ sites.

To examine which lysine residue(s) may be the main ubiquitylation sites, we generated G6PD mutants bearing single or double lysine-to-arginine (Lys-to-Arg) substitutions and polyubiquitylated G6PD was assessed. As shown in Figure 9B and 9C, compared with wild-type G6PD, Ub conjugation to G6PD mutants with either Lys366-to-Arg (K^366R^) substitution or Lys403-to-Arg (K^403R^) substitution was largely reduced. However, the ubiquitylation of the other G6PD mutants (K^97R^, K^407R^ and K^408R^) was not significantly affected. Remarkably, Lys-to-Arg substitutions on both K^366^ and K^403^ sites (K^366+403R^) almost completely abolished the VHL-mediated ubiquitylation on G6PD (Figure 9C).

## Discussion

The current study reveals the biochemical mechanism that high glucose promotes the degradation of G6PD protein through the ubiquitin proteasome pathway, which leads to podocyte injury. Further, VHL, an E3 enzyme subunit, plays a key role in the ubiquitylation and degradation of G6PD, which is an important anti-oxidant component.

The injury and loss of podocytes plays a critical role in the initiation and development of DKD, while the mechanisms are not fully revealed[31]. Here, our results show that G6PD protein level in both podocytes from diabetic patients and podocytes exposed to high glucose are significantly decreased, which may explain the injury and loss of podocytes in DKD.

G6PD is the rate-limiting enzyme in the pentose phosphate pathway, which plays a critical role in cell growth by providing NADPH for redox regulation[32, 33]. Cells with normal G6PD activity keep the net ROS production at a reasonably low level. However, G6PD deficiency will cripple the antioxidant defense, resulting in the build-up of oxidative damage. Previously, we and others showed that G6PD deficiency led to increased accumulation of ROS in many cell types, which impaired the cellular function and survival[18, 21]. In this study, we reported, for the first time, that high glucose-induced G6PD deficiency in podocytes resulted in increased ROS accumulation and apoptosis, which could be rescued by overexpressing G6PD. We previously demonstrated that non-diabetic G6PD deficient mice had increased urinary albuminuria[22] and here in the same mouse model, we reported that G6PD deficiency led to significant podocyte loss due to increased apoptosis. Since podocytes are terminally differentiated epithelial cells, the loss of podocytes would eventually lead to the increased urinary albuminuria. The above results demonstrated that G6PD played vital roles in maintaining the survival and function of podocyte.

Due to the decrease of G6PD induced by high glucose, NADPH and GSH levels were decreased, which resulted in the depletion of glutathione stores and enhanced oxidative stress. Further support for the role of G6PD in regulation of NADPH and GSH was that overexpression of G6PD conferred protection on podocytes exposed to high glucose. There is accumulating evidence to support this finding in other cell types. Mouse embryonic fibroblasts cells with increased G6PD level are more resistant to the oxidant tert-butyl hydroperoxide than cells with low G6PD activity[34]. Our previous studies have shown overexpressing G6PD prevents the high glucose-mediated ROS accumulation in both endothelial cell and pancreatic β cells[18, 21]. As for the antioxidant defense of podocytes, previous studies have proven that leukemia inhibitory factor protects against the high glucose-induced podocyte apoptosis through inhibiting oxidative stress[35]. Inhibiting the epidermal growth factor receptor in podocytes can decrease high glucose-induced ROS production[36]. Here, we demonstrate that in podocytes, G6PD is also critical in promoting cellular resistance to oxidative stress induced by hyperglycemia.

Work from others have reported that G6PD expression was up-regulated in pancreatic islets, adipose tissue and liver of diabetic animals and the over-activation of G6PD would stimulate ROS production[37–40]. The disparity between their work and our findings suggest that the expression of G6PD in various tissues may respond to hyperglycemia differently and either G6PD over-activation or deficiency would induce the accumulation of ROS by distinct mechanisms depending on the cell type.

G6PD is subject to complex regulation, and modifications of G6PD protein have been reported[15, 41, 42]. It has been proven that G6PD acetylation on K^403^ affects the formation of active dimers, which decreases G6PD activity[15]. Conversely, two other studies have reported that glycosylation on serine 84 and sirtuin 5-associated deglutarylation of G6PD increase G6PD activity[41, 42]. In this study, we present a novel finding for the regulation of G6PD protein. We verified that K^366^ and K^403^ in G6PD were the major sites for VHL-mediated ubiquitylation. The ubiquitylation of G6PD on K^97^ was also suggested in previous studies based on mass spectrometry analysis, while no functional validation was performed[30, 43]. Work from us and Wang *et al* showed that the modifications on K^403^ decreased G6PD activity through different mechanisms. We revealed that ubiquitylation on K^403^ promoted G6PD protein degradation, while Wang *et al* validated that acetylation on K^403^ impaired the formation of active dimer. This further suggests that K^403^, an evolutionarily conserved residue, is very critical for maintaining the spatial conformation and protein stabilization of G6PD.

VHL gene is on the short arm of chromosome 3 (3p25-26)[44]. Through forming the VCB-Cul2 complex including elongin B, elongin C, and Cullin 2, the VHL protein is part of an E3 ubiquitin ligase[45]. Inactivation of VHL is associated with several tumors, such as sporadic renal clear cell carcinomas and pancreatic neuroendocrine tumors[46, 47]. In addition, studies have reported that specific deletion of VHL in pancreatic β cell results in impaired glucose stimulated insulin secretion, indicating that VHL may participate in the pathogenesis of diabetes[48–50]. Additionally, it was observed that podocyte-specific VHL knockout led to rapid progressive glomerulonephritis, which was attributed to the increased hypoxia-inducible transcription factor-1α, the most well-known target protein of VHL[51]. There were 5 other identified VHL target proteins, including extracellular signal-regulated kinase 5[52], sprouty2[53], β_2_-adrenergic receptor[54], IκB kinase-β[55] and ceramide kinase like protein[56]. However, the potential role of VHL in the pathogenesis of diabetic kidney disease has not been determined. Here, for the first time, we show that G6PD is a novel target protein of VHL and VHL / G6PD axis plays an important role in maintaining the function and survival of podocytes in DKD. Taken together, it is enticing to further explore the potential role of VHL / G6PD as a new therapeutic target for diabetic kidney disease.

## Materials and Methods

### Human renal biopsy samples

Renal tissue samples were obtained from 3 diabetic patients who underwent renal biopsy in Division of Nephrology of Huashan Hospital and had been confirmed to have pathological and clinical findings consistent with diabetic kidney disease. All participants provided written informed consent, which was approved by the ethics committee at Huashan Hospital (KY2016-394). Normal human kidney tissues (n=3) without diabetes or renal disease were obtained via autopsy.

### Animal study

The G6PD-deficient mouse model was recovered in the offspring of 1-ethyl-nitrosourea-treated male mice on a C3H murine background by Pretsch *et al*[26]. Later, Sanders *et al* showed that there was a single-point mutation (A to T transversion) in the 5’ splice site consensus sequence at the 3’ end of the exon 1[57]. The mice were bred at Brigham and Women’s Hospital and Harvard Medical School from frozen embryos obtained from the Medical Research Council. This animal model was previously characterized, and mice were genotyped by polymerase chain reaction as described previously[58]. Hemizygous (Hemi) G6PD-deficient male mice, which had 15% of wild type G6PD activity, and age-matched wild type C3H control mice, aged 9wk, 23wk and 39wk, respectively, were used.

Male Sprague-Dawley rats (SLRC Laboratory Animal) weighing 240-260g were maintained on a standard chow with free access to water. Rats were randomly divided into control and diabetic groups. Diabetes was induced by an intraperitoneal injection (ip) of streptozotocin (STZ) (Sigma) in citrate buffer (0.1mol/l, pH4.5) with a dose of 55mg/kg body weight. The non-diabetic (NDM) control group received injection of citrate buffer alone. Blood glucose level was measured 7 days after STZ injection, and the rats with blood glucose level higher than 16.7mmol/L were considered diabetic. Rats were sacrificed 12 weeks after the onset of diabetes.

5-week-old male DBA/2J mice (SLRC Laboratory Animal) were made diabetic after injected with STZ (50mg/kg, ip) on 5 consecutive days. Hyperglycemia was confirmed when the blood glucose level reached 16.7mmol/L at 1-week post-injection. Mice were sacrificed 12 weeks after the onset of diabetes.

Male DBA/2J genetic background Akita mice bearing Ins2+/C96Y mutation[59] from The Jackson Laboratory were maintained on standard chow and had free access to water. Wild type littermates were used as non-diabetic controls. Mice were sacrificed at 6 months old.

All animal experiments were approved by the Institutional Animal Care and Use Committee at Harvard Medical School or Fudan University and conducted in accordance with the Guide for the Care and Use of Laboratory Animals published by the US National Institutes of Health (NIH Publication No. 85-23, revised 1996).

### Cell culture

Conditionally immortalized mouse podocytes were cultured as described previously[60]. Podocytes were exposed to RPMI 1640 (Gibco) medium containing different concentrations of either glucose (5.6mM, 25mM or 5.6mM supplemented with 19.4mM mannitol) or palmitate (125μM, 250μM or 500μM).

Human embryonic kidney (HEK293T) cells were maintained in Dulbecco’s Modified Eagle Medium (Gibco) supplemented with 10% fetal bovine serum.

### siRNA transfection

Cells were seeded into 6-well plates and grown until 60-80% confluent. siRNA for G6PD or VHL was transfected with lipofectamine TM RNAiMAX Transfection Reagent (Invitrogen) following the recommended protocol.

The sequences designed for inhibiting G6PD gene expression were 5′-AAUCAACUGUCGAACCACAtt-3′ and 3’-UGUGGUUCGACAGUUGAUUgg-5’.

A scrambled siRNA (4390846, Invitrogen) without biological effects was used as control.

The sequences designed for inhibiting VHL gene expression were listed below:
siVHL#1 were 5′-GCUCUACGAAGAU CUGGAAdT dT-3′ and 3’-UUCCAGAUCUUCGUAGAGCdT dT-5’; siVHL#2 were 5′-GCAUUGCACAUCAACGGAUdTdT-3′ and 3’-AUCCGUUGAUGUGCAAUGCdTdT-5’.

The control siRNA for VHL was obtained from Biotend (N100, Biotend).

### Adenoviral vector construction

Human G6PD cDNA was excised from pCMV-XL4-G6PD (OriGene) and was confirmed by sequencing. The adenoviral-hG6PD expression vector was constructed as described previously[21]. MOI 10 was used for all experiments. Empty vector was used for control experiments.

### Real-time PCR analysis

Real-time PCR was performed as described before[61]. Total RNA was extracted from podocytes with TRIzol Reagent (Invitrogen) and was converted to cDNA according to the manufacturer’s protocol (G490, ABM). The sequences of the primers used were listed below: mouse G6PD were 5’-CAGCGGCAACTAAACTCAGAA-3’ and 3’-GCATAGCCCACAATGAAGGT-5’; mouse β-actin were 5’-CACGATGGAGGGGCCGGACTCATC-3’ and 3’-TAAAGACCTCTATGCCAACACAGT-5’.

### Western Blot

Cells were lysed with lysis buffer containing protease inhibitor cocktail (Roche). The membranes were incubated with antibodies to G6PD (Bethyl Laboratories), cleaved caspase-3 (Cell Signaling Technology), β-actin (Santa Cruz Biotechnology), GAPDH (Santa Cruz Biotechnology), anti-HA antibody (Sigma), anti-Flag antibody (Sigma) and anti-His antibody (Cell Signaling Technology).

### Measurement of oxidative stress markers

Measurement of NADPH, GSH/GSSG and ROS was performed as described before[18, 21]. NADPH was measured by a colorimetric method according to the manufacturer’s instructions (Bioassay System). GSH/GSSG was measured by a spectrophotometric method according to the manufacturer’s instructions (Cayman). ROS production was measured with the cell-permeable, oxidation-sensitive dye CM-H_2_DCFDA (Invitrogen). Fluorescence was read with a microplate fluorometer (Victor2 fluorometer, PerkinElmer). After reading, cells were lysed to measure protein concentration, which was used to normalize the readings.

### Immunohistochemistry

Expression of WT-1 was examined on paraffin-embedded renal tissue section with 1:100 dilution of WT-1 antibody (Santa Cruz Biotechnology). The number of WT-1 positive stained cells per glomerular cross-section area in the kidney sections was analyzed as described previously[62].

### Immunofluorescence

Immunofluorescence staining of paraffin blocks was applied to assess the expression of WT-1 and G6PD in renal tissue section with WT-1 antibody (Santa Cruz Biotechnology, 1:100) and G6PD antibody (Proteintech Group, 1:200).

### TUNEL analysis

The terminal deoxynucleotidyl transferase-mediated dUTP nick end–labeling (TUNEL) staining was carried out on 4-mm-thick paraffin–embedded sections using a cell death detection assay kit according to the manufacturer’s instructions (Roche). Samples were evaluated under a Nikon Eclipse ci-L fluorescence microscope (Nikon, Japan).

### Yeast two hybrid screen

The yeast two hybrid (Y2H) screening was carried out to screen for E3 ubiquitin ligases that may ubiquitylate G6PD. We utilized the GAL4-based yeast two-hybrid system (Y2H, Invitrogen) to screen for and analyze the protein-protein interaction in yeast as described before[63]. The full length of G6PD open reading frame was first cloned in donor vector pDONR221 and then transferred into pDEST32 through Gateway cloning reaction (Invitrogen), generating the bait plasmid, pDEST32-G6PD, which contained the in-frame fusion of GAL4 DNA binding domain. The prey vector pDEST22 containing human cDNA collections in-frame fused to the GAL4 activating domain (Invitrogen). Using the empty pDEST22 plasmid as a negative prey control, Y2H screening was performed by transforming yeast strain (Mav203 strain) that harbored bait vector, pDEST32-G6PD, with the prey vectors for human E3 cDNA expression library. Yeast transformants were first grown on to the agar plate on SD-2 (deficient in Leucine and Tryptophan) for selecting yeast cells containing both bait and prey vectors, and then transferred to SD-4 (deficient in Leucine, Tryptophan, Histidine, and Uracil) plates to screen for proteins that potentially interacted with human G6PD. Colonies grown on the SD-4 plates were picked and streaked onto another SD-4 plates with X-Gal (5-Bromo-4-chloro-3-indolyl β-D-galactopyranoside, Sigma) added. “Positive” colonies were scored for those which not only grew in SD-4 medium but also presented blue color in X-Gal staining assay for β-galactosidase activity. The prey vectors were recovered from the positive colonies and sequenced after amplification in *E. coli.* Each interaction was confirmed by transforming yeast Mav203 cells with the indicated bait and prey vectors. The transformants grew on the SD-2 or SD-4 agar plates (with or without X-Gal) for approximately 3 days at 30 °C. Images of the colonies on both plates were recorded.

### Immunoprecipitation (IP) and immunoblotting (IB)

HEK293T cells were transiently transfected with indicated expression plasmids with lipofectamine™ 2000 Transfection Reagent (Invitrogen). Usually, cells were harvested 48 hours after transfection and washed twice with ice-cold PBS buffer. Cells were then sonicated in IP buffer [20mM Tris-Cl, 150mM NaCl, 1mM EDTA, 1mM EGTA, 1% (v/v) Triton X-100, 2.5mM sodium pyrophosphate, 1mM β-glycerolphosphate, 1mM Na_3_VO_4_, and protease inhibitor cocktail (Roche), pH7.5] by Bioruptor UCD-200 (Diagenode) and then centrifuged at 22,500g at 4°C for 15 min. Expression of the indicated proteins in the lysates was checked by immunoblotting with relevant antibodies to normalize total input amounts. After normalization, the supernatants were each incubated with equal amounts of anti-Flag M2 beads overnight at 4°C. The anti-Flag M2 beads and interacting proteins were pelleted and washed three times with IP buffer before boiling in 1X SDS-PAGE sample. The boiled samples were then resolved in SDS-PAGE and subjected to immunoblotting analysis with indicated antibodies.

### In vivo ubiquitylation assay

To check the ubiquitylation status of G6PD, Flag-G6PD was immunoprecipitated from the cells treated with proteasome inhibitor MG132. For Figure 7E and Figure 8E, HEK293T cells were transiently transfected with indicated expression plasmids with lipofectamine™ 2000 Transfection Reagent (Invitrogen). After cells were harvested, IP experiments were carried out in RIPA buffer [50mM Tris-Cl, pH7.4, 150mM NaCl, 5mM EDTA, 1% (v/v) Triton X-100, 0.5% sodium pyrophosphate, 0.1% SDS, and protease inhibitor cocktail (Roche)]. The cell lysates were centrifuged at 22, 500g at 4°C for 15 min. The supernatants were subjected to immunoblotting to confirm the expression of each protein or incubation with anti-Flag M2 beads overnight at 4°C. The recovered beads were then washed three times and finally boiled in 1X SDS-PAGE sample loading buffer, followed by SDS-PAGE and immunoblotted using anti-His.

### Mass Spectrometry Analysis

Samples were prepared with the same protocols as described at in vivo ubiquitylation assay. After the supernatants were incubated with anti-Flag M2 beads overnight at 4°C and washed three times, the protein was eluted by Flag peptide (ApexBio technology). Sample analysis was performed on nano-scale HLPC-MS system as described previously[63].

### Statistical analyses

All data were expressed as mean±sd from three independent experiments. Statistical analysis was performed with a two-tailed unpaired Student’s t-test. The *P* value less than 0.05 was considered statistically significant.

## Acknowledgements

We thank Zhihong Yang at Joslin Diabetes Center for supplying podocytes to us. Thank Min Zhang in Division of Nephrology of Huashan Hospital for cell culturing guidance. Thank Wei Huang at Fudan University for helpful comments with the manuscript. We also acknowledge the excellent support from all members of Ronggui Hu’s laboratory. In addition, this study was supported by the grants from National Natural Science Foundation of China (no.81370938, no.81471041 and no.81400796).

## Author contributions

ZYZ conceived the original idea of this study. JH, RCS and ZYZ designed experiments. MW, LLY, YPY, SZG, MW and YY collected samples and performed experiments. MW, JH, MH, WJL, QL, WG, YA, BL, CMH, QHW, YML, RGH, RCS and ZYZ analyzed and interpreted the data. DEH provided mouse model. MW, RCS, QHW, DEH and ZYZ prepared the manuscript with suggestions from all other authors.

## Conflict of interest

All the authors declared no competing interests.

---

**Figure EV1.**
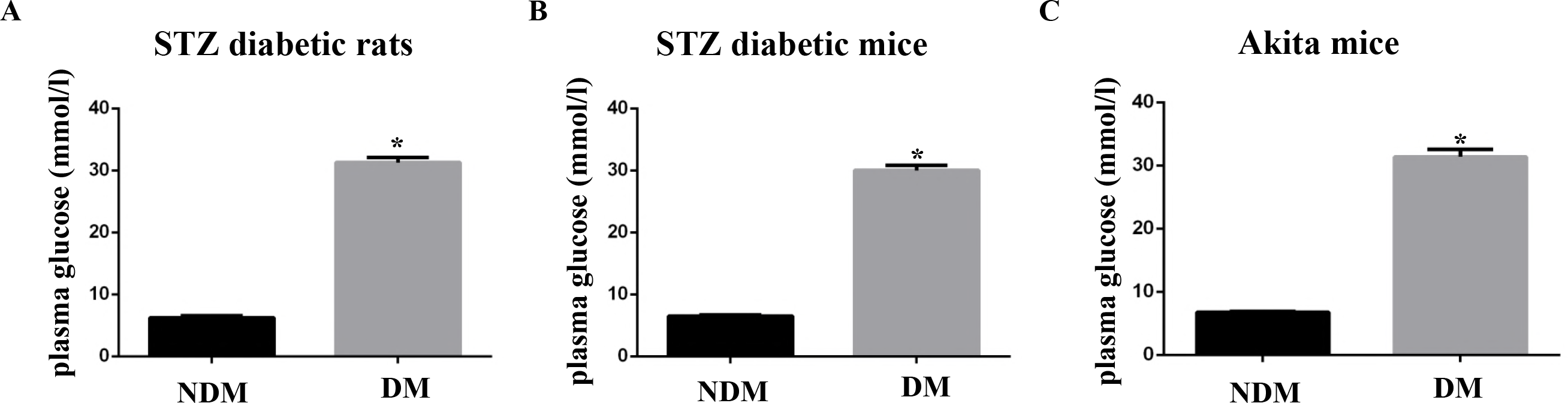
Blood glucose was increased in diabetic rodents. A, B, C Blood glucose was increased in STZ-induced diabetic rats, STZ-induced diabetic mice and Akita mice, respectively. n=6 mice for each group. ^*^ denotes *P* < 0.05 for DM versus NDM.

**Figure EV2.**
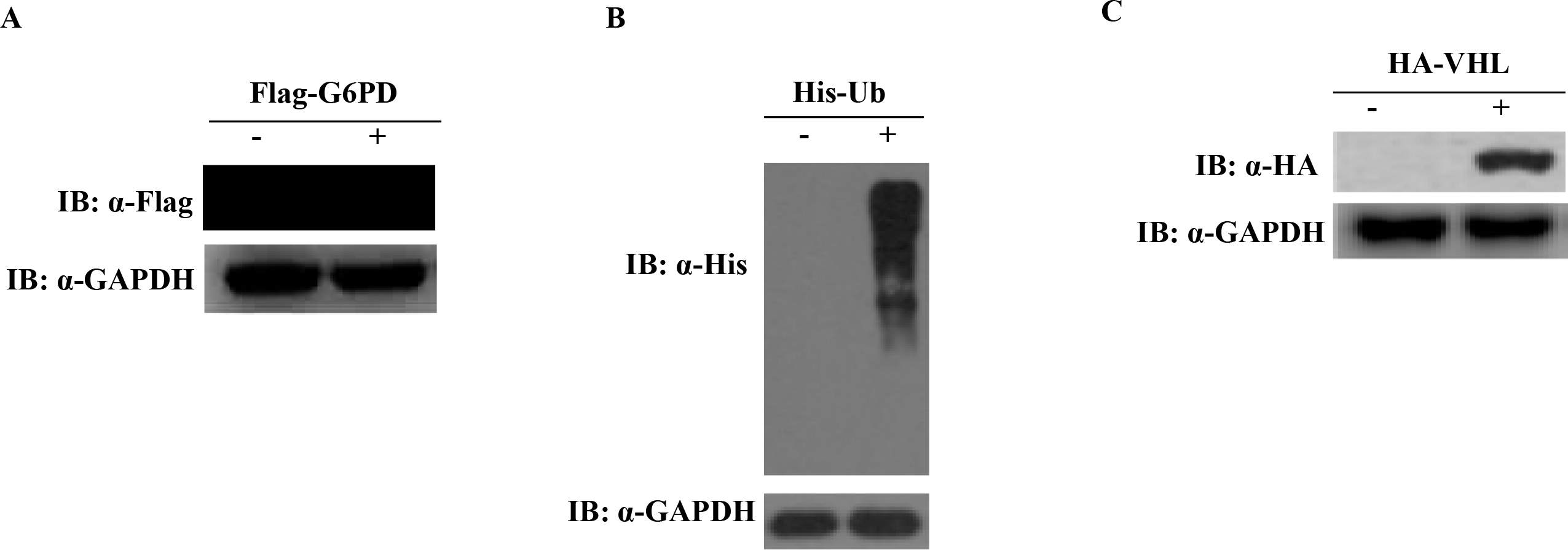
Verification of Flag-G6PD, His-Ub and HA-VHL plasmids. A Flag-G6PD plasmid was expressed in HEK293T cells. Flag-G6PD plasmid was transfected into HEK293T cells for 48 hours. Cells lysates were subjected to Flag and GAPDH immunoblotting. B His-Ub plasmid was verified in HEK293T cells. HEK293T cells were transfected with His-Ub plasmid for 48 hours. Protein level of His was analyzed by Western blot with anti-His antibody. C HA-VHL plasmid was expressed in HEK293T. HEK293T cells were transfected with HA-VHL plasmid for 48 hours and cells lysates were subjected to HA and GAPDH immunoblotting.

